# Cortico-hippocampal morphology and behavioural indices improved in maternal deprivation model of schizophrenia following vitamin B complex supplementation

**DOI:** 10.1101/2020.06.16.154468

**Authors:** Gabriel Olaiya Omotoso, Fatimah Adeola Abdulsalam, Nafisat Yetunde Mutholib, Abdulkabir Bature, Ismail Temitayo Gbadamosi

**Author notes:** **Corresponding Author:** Dr G. O. Omotoso, Department of Anatomy, Faculty of Basic Medical Sciences, College of Health Sciences, University of Ilorin, P.M.B. 1515, Ilorin, 240003, Nigeria.

## Abstract

Maternal deprivation (MD) during early life development has been documented to culminate in long-term alterations in brain function and behavioural manifestations that mimic schizophrenia. This study elucidated the putative neuroprotective roles of vitamin B complex in MD-induced behavioural and neurochemical modifications in hippocampus and prefrontal cortex of Wistar rats. Rat pups were maternally deprived on postnatal day 9 for 24 hours and then treated with or without vitamin B complex for 15 days while a control group was undisturbed during the experimental period. The rats were then subjected to behavioural paradigms to measure memory indices and anxiety levels. The rats were sacrificed to obtain the PFC and hippocampus for histomorphological and biochemical analysis. Behavioural analysis of the animals revealed that MD induced a declination in long- and short-term memory in addition to anxiety-like behaviour in the open field test. Cortico-hippocampal histomorphology of these animals showed an increased astrocytic density and chromatolysis, which were accompanied by reduced levels of superoxide dismutase and catalase enzymes. Vitamin B complex mitigated MD-induced behavioural decline, histomorphological perturbation and oxidative stress by enhancing the intrinsic antioxidant defence, thereby culminating in nootropic behaviour and reduced anxiety. In conclusion, we confirmed the hypothesis that vitamin B complex is neuroprotective against neuropathological alterations induced by maternal deprivation.

## 1. Introduction

Neurodevelopment is a function of genetic and environmental interactions (Crawford et al., 1993). Proper nutrition, being an environmental factor, is essential during the critical periods of neurodevelopment (Casper, 2004). The development of cognitive, motor, and socio-emotional skills throughout adolescence and adulthood are established during this critical period of life (Prado and Dewey, 2014). Breastfeeding is a vital aspect of growth and development because a developing mammal derives a substantial amount of nutrient required for neurodevelopment from the breast milk of its mother during the initial phase of postnatal life (Crawford et al., 1993). Events in early life are important in determining normal physiological functions in later life, and predisposition towards pathophysiological processes, including behavioural imbalances or inappropriate stress responses (Viveros et al., 2009).

New-born mammals of many species naturally display a high dependence on maternal care during the first days or weeks of life, making this time a period of extreme susceptibility to changes in mother-infant interactions (Lehmann and Feldon, 2000). Breastfeeding is believed to be linked with healthier neurologic outcomes from infancy to adulthood. In most species, the mother provides not only breast milk but also triggers the pup’s normal development and provides warmth as well as protection for the new-born. Consequently, the mother-pup contact is important for survival and normal development of the new-born (Lehmann and Feldon, 2000). Animal studies and human data have proposed that the relationship between the quality of early environmental and emotional responses during adulthood appears to be mediated by parental/maternal influences on brain development (Llorente et al., 2010; Petryk et al., 2007).

Maternal separation and nutritional deficiency are risk factors for developing schizophrenia (Walker and Diforio, 1997), a neurological condition that affects about 1% of the global population (Andreasen et al., 2002; Schoenrock and Tarantino, 2016). Due to non-availability of a definitive therapeutic cure, combination therapy is usually adopted, involving psychotherapy, pharmacotherapy including use of atypical and typical antipsychotics, and other social interventions (Kapur and Remington, 2001; Padurariu et al., 2010; Raffa et al., 2009).

Although the main pathogenesis of the disorder is unknown, oxidative stress is pivotal in the pathophysiology of schizophrenia (Yao et al., 2001). The brain is more vulnerable to oxidative stress due to its high oxygen consumption, compared to other body organs (Bitanihirwe and Woo, 2011). While it is common for schizophrenic patients to have low levels of antioxidant enzymes, many studies have reported inconsistent data (Bitanihirwe and Woo, 2011; Padurariu et al., 2010; Zhang et al., 2010). Hence the need to know the factors that determine the level of each antioxidant enzyme during oxidative stress associated with schizophrenia and how such enzyme markers can be effectively utilized as diagnostic tools. There are contradictory results of antioxidant enzymes such as superoxide dismutase, catalase, and glutathione peroxidase in schizophrenic patients with different authors reporting either reduced or increased level of the same enzyme in some cases, or that the activity of a particular antioxidant enzyme was not affected by this condition (Padurariu et al., 2010; Zhang et al., 2010).

Dietary vitamin supplementation is used in the management of several clinical conditions (Delanty and Dichter, 2000; Schoenrock and Tarantino, 2016; Zugno et al., 2015). The B-vitamins are a group of water-soluble vitamins that play vital roles in brain development and ultimately maintaining cognitive functions (Stough et al., 2014). They are involved in energy production, DNA synthesis, and DNA repair, and these underscore their usefulness in the prevention and amelioration of neurological diseases and disability (Ford et al., 2018; Reynolds, 2006). In a human population-based study by Cao *et al*., pyridoxine and nicotinamide were found to show strong association with schizophrenia (Cao et al., 2018). A study observed that mega doses of some B-vitamins reduced pathologies associated with schizophrenia (Hoffer, 1971). As a result of the possibility of a synergy of functions, it is rational to use the B vitamins in combination, rather than a therapeutic use of individual members of the B complex (Kennedy, 2016).

This study aimed to characterize the behavioural, histomorphological and neurochemical changes in the prefrontal cortex and hippocampus of maternally deprived Wistar rats and subsequently elucidating the role of supplementation with vitamin B complex.

## 2. Materials and methods

### 2.1 Ethical approval

This study was approved by the University of Ilorin Ethical Review Committee. The procedures and experiments in this study were conducted with strict compliance with the Institutional Animal Care and Use Committee guidelines.

### 2.2 Experimental animals

Ten adult male and twenty female Wistar rats were obtained and used for the study. They were housed and acclimatized in the Animal House Facility of the Faculty of Basic Medical Sciences, University of Ilorin, Ilorin, Nigeria. They had liberal access to rat chow and water. The oestrous cycle of the female rats was determined following the vaginal smear method of Marcondes et al., 2002. The rats were mated by housing one male rat with two female rats during the oestrous phase. After the pregnant rats delivered, litters were culled to nine (9) pups per dam. The offspring were weaned at PND 21.

### 2.3 Experimental design

The pups were divided into four groups A-D. Group A served as the control group, Group B was treated with vitamin B complex from postnatal day (PND) 21-35, Groups C and D were maternally deprived for 24 hrs on PND 9, Group D was subsequently treated with Vitamin B complex from PND 21 to 35.

### 2.4 Maternal deprivation protocol

We adopted the protocol described by Llorente *et al*. (2007). On the first postnatal day, the litters were reduced to nine pups per group. The pups were deprived of their mother for 24 h on PND 9 beginning at 9:00 am by removing the mother from the cage and maintaining the pups in the cage in the presence of the maternal odour. On PND 10, the pups were weighed, and the mothers were returned to the pups.

### 2.5 Vitamin Supplementation

Vitamin B complex tablets were procured from Prime Health Pharmacy, Ilorin. The four components of the complex were: Vitamins B_1_, B_2_, B_3_ and B_6_ with 5 mg, 2 mg, 20 mg, and 2 mg composition respectively, administered orally at a daily dosage of 40 mg/kg (Khan *et al*., 2008), and treatment commenced from PND 21 to 35.

### 2.6 Neurobehavioural studies

At the end of treatment, the animals were subjected to Morris water maze, Y-maze, and open field behavioural paradigms to measure the animals’ spatial memory index, working memory index, and anxiety levels.

#### 2.6.1 Open field test

The open field test was carried out as previously described in literature (Gould et al., 2009). The open-field device consisted of plywood measuring 100 cm x 100 cm with walls 50 cm high. The floor was sectioned into square grids and a centre square. During the test, the rats were placed in the centre square and allowed to explore for 5 minutes. The activities of the animals were recorded by a video hanging from above. Five behaviors were scored, and these included the number of lines crossed, centre square entry, centre square duration, rearing frequency and stretch attend posture. The number of lines crossed was the frequency with which the rats crossed one of the grid lines with all four paws. The number of liners and rearing frequency was deduced from the activities of the rats in the open field.

#### 2.6.2 Y-maze spontaneous alternation test

The test was conducted as described by Wright *et al*. (2006), with slight modification. It assessed hippocampal-dependent spatial recognition memory. The Y-maze consisted of three identical wooden arms (50 cm long, 16 cm wide and 32 cm high) and floor of each arm was made of plywood. The rats were placed in the centre of the maze ad allowed to explore for five minutes. The sequence of alternation was recorded and analyzed to obtain the percentage correct alternation (Wright et al., 2006).

#### 2.6.3 Morris water maze test

The Morris water maze is used to quantify spatial memory and learning (Nunez, 2008). The water maze consisted of a round pool about six feet in diameter and about 3 feet deep. The rats were placed in a small pool of water containing an escape platform placed an inch deep to the surface of the water. The animals were trained 24 hrs prior to the main test day. During the test, the animals were placed in the pool of water and allowed to find the escape platform themselves or by guidance; this was repeated three times. On the test day, the water was coloured, and the animals were allowed a maximum period of 60 seconds to find the escape platform. The time taken to find the escape platform is recorded as the escape latency period.

### 2.7 Animal sacrifice for histological demonstration

Rats for histological and histochemical studies were euthanized and subjected to transcardial perfusion using normal saline, followed by 4% paraformaldehyde (PFA). The brain tissues were then excised, weighed and post-fixed in 4% paraformaldehyde. The tissues were processed for histological and histochemical examination and stained using hematoxylin and eosin (Fischer et al., 2008) and cresyl fast violet (for Nissl bodies demonstration). Immunohistochemical study was conducted to examine the expression of astrocytes using anti-glial fibrillary acidic protein antibody (Goldstein and Watkins, 2008). This involved an immunoperoxidase technique comprising the processes of heat-induced antigen retrieval with citric acid, endogenous peroxidase blocking using hydrogen peroxide, 10% calf serum with 1% BSA and 0.1% Triton X-100 as blocking buffer, primary antibody (anti-GFAP) and secondary antibody (horseradish peroxidase); chromogen 3.3□-diaminobenzidine for colour intensification, and finally counterstained with hematoxylin. Stained slides were dehydrated, cleared, and mounted.

### 2.8 Animal sacrifice for biochemical studies

Rats for enzymatic studies were sacrificed by cervical dislocation; the brains were then removed, weighed and rinsed in 0.25 M sucrose each and stored in 30% sucrose at 4°C. The hippocampus and prefrontal cortex were excised, weighed and homogenized in 0.25 M sucrose at 4°C. The tissue homogenate was centrifuged for 15 min in a centrifuge at 5000 rpm to obtain supernatants which were aspirated into plain labeled glass cuvettes placed in ice. Biochemical assay for superoxide dismutase (Cat #: KT-60703), glutathione peroxidase (Cat #: MBS744364), catalase (Cat #: MBS701713) and malondialdehyde (Cat #: MBS9389391) levels were carried out on the supernatants obtained using spectrophotometric techniques according to manufacturer’s instruction in the assay kit pack.

### 2.9 Photomicrography and Statistical analysis

Histology and IHC images were acquired using an Olympus binocular research microscope (Olympus, New Jersey, USA) connected to an Amscope Camera (5.0 MP). All quantitative data were analyzed using GraphPad® (version 7) software. The results of the body weight, number of lines crossed, rearing frequency and oxidative markers were analysed using oneway ANOVA with Tukey’s multiple comparisons test. Significance was set at p values less than 0.05.

## 3 Results

### 3.1 Changes in body and brain weights

A decrease in body weight was seen in all treated groups compared to the Control, with maternally deprived rats having the lowest weight gain, as revealed by the low weight difference. Maternally deprived rats that received vitamin B complex intervention had a higher weight gain compared to maternally deprived rats that did not receive vitamin B complex, although the former was still lesser than the body weight of vitamin B complex-treated rats. There was no significant difference in the brain weight of the treated groups compared to control, though maternally deprived rats had the least brain weight. Meanwhile, maternally deprived rats that were treated with vitamin B complex had a significant increase in brain weight compared to those that did not receive vitamin B complex (p<0.05) (Table 1). Although the weight of the prefrontal cortex (Table 1) was lowest in maternally deprived rats compared with the Control and other treated groups, this was not statistically significant (p>0.05). Meanwhile, maternally deprived rats that received vitamin B complex had an increase in PFC weight compared to those that received no vitamin B complex. Weight change in the hippocampus (Table 1) was more remarkable compared to the prefrontal cortex, as a significant decrease was seen in the maternally deprived group compared to control (p<0.05) and vitamin B complex-treated rats (p<0.01); however, intervention with vitamin B complex improved hippocampal weight.

**Table 1:** Body and brain weights (n=5 per group)

| Groups | Initial Body weight (g) | Final Body weight (g) | Weight Difference (g) | Brain weight (g) | Relative Brain weight (g) |
| --- | --- | --- | --- | --- | --- |
| **Control** | 10.46±0.26 | 40.76±1.30 | 30.3 | 1.23±0.12 | 0.0302 |
| **VIT BCO** | 10.73±0.63 | 39.33±3.07 | 28.6 | 1.19±0.12 | 0.0303 |
| **MD** | 10.66±1.44 | 34.86±0.51 | 24.2 | 1.11±0.06 | 0.0318 |
| **MD+VIT BCO** | 10.01±0.87 | 37.08±1.04 | 27.1 | 1.37±0.04* | 0.0369 |

VIT BCO: vitamin B complex-treated; MD: maternally deprived; \* significant increase when compared with MD rats ( $p<0.05$ ).

### 3.2 Vitamin B complex attenuated MD-induced behavioural perturbation

The open-field behavioural paradigm was used to access the anxiety-like behaviour of the experimental animals in this study. There was a significant reduction in the rearing frequency in maternally deprived rats compared to rats that received vitamin B complex (p<0.05) and control (p<0.05) group (Fig. 1a). Likewise, a significant reduction in the number of lines crossed in maternal deprivation rats compared to rats that received vitamin B complex (p<0.05) and control (p<0.05) group (Fig. 1b) was also observed. Oral administration of vitamin B complex prevented this anxiety-like behavioural changes in rats that were maternally deprived then treated with vitamin B complex, as it was observed in the number of lines crossed which was significantly higher than those of maternally deprived group (p<0.05).

**Figure 1:**
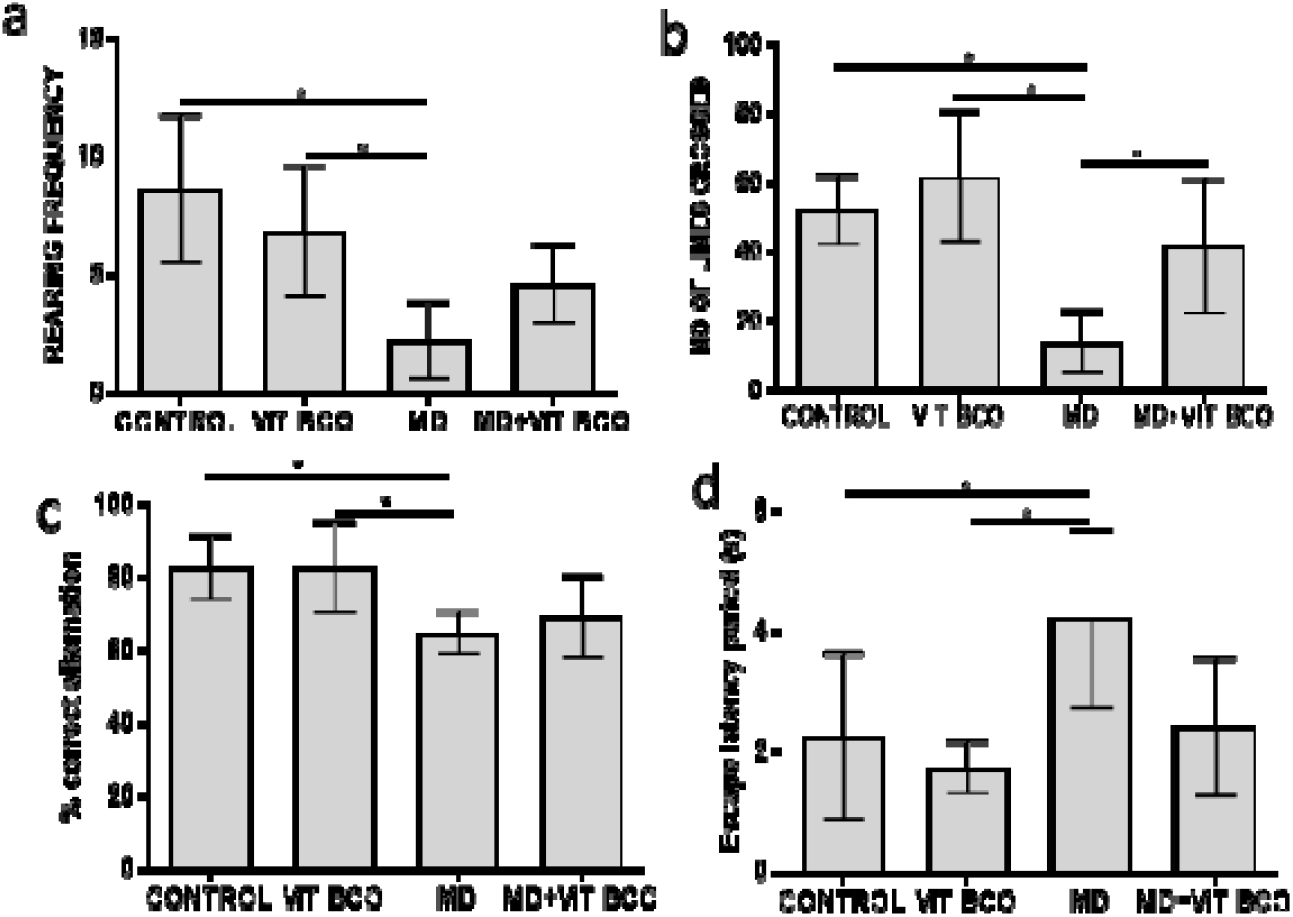
Behaviour; rearing frequency (a) and number of lines crossed (b) of experimental animals in the open-field behavioural paradigm. Percentage correct alternation of rats in Y-maze (c) and Escape latency period of experimental animals in the Morris water maze (d). The rearing frequency of MD group was markedly lower than those of the control (p<0.05) and VIT BCO group (p<0.05). MD groups presented with a significantly lower number of lines crossed relative to other experimental groups. MD group showed a significantly lower % correct alternation relative to the control (p<0.05) and VIT BCO group (p<0.05). Escape latency of MD group was significantly higher relative to control (p<0.05) and VIT BCO (p<0.05) group. VIT BCO= Vitamin B complex, MD= maternal deprivation, MD + VIT BCO = maternal deprivation + Vitamin B complex. * is p-value <0.05.

In the Y-maze behavioural paradigm, the percentage correct alternation was used to quantify short-term memory. Maternal deprivation caused a reduction in percentage correct alternation in rats compared to rats that received vitamin B complex (p<0.05) and control (p<0.05) group (Fig. 1c). Maternally deprived rats presented with significant elevated escape latency time in Morris water maze relative to rats that received vitamin B complex (p<0.05) (Fig. 1d). However, vitamin B complex prevented the reduced short-term memory indices in maternally deprived rats treated with vitamin B complex. This finding was corroborated with the findings from the Morris water maze. The escape latency time in Morris water maze was used to measure long term memory index in animal studies.

Cumulatively, this result from the behavioural analysis of the animals in the open field test, Y-maze, and Morris water maze points out that maternal deprivation mediated manifestation of anxiety and reduced memory performance both of which were attenuated by vitamin B complex.

### 3.3 Vitamin B complex ameliorated MD-induced oxidative stress

The activities of antioxidant enzymes and lipid peroxidation were quantified to elucidate the latent effect of MD and VIT BCO on neural oxidative status. Superoxide dismutase (SOD) activity (Fig. 2a and b) was significantly reduced in the MD group relative to the control group (p<0.05 for PFC and p>0.05 for hippocampus). This finding suggests that SOD is the main antioxidant enzyme affected by MD. SOD plays a pivotal role in the frontline of the antioxidant defence system where it catalyses the breakdown of highly reactive superoxide anions to less reactive hydrogen peroxide. Hence, a depleted level of neural SOD will result in accumulation of superoxide anions that will drive lipid peroxidation. Glutathione peroxidase (GPX) activities in both brain regions showed no significant variation across the experimental group (Fig. 2c and d). MD group presented with a reduction in the level of catalase in the PFC (p>0.05) and hippocampus (p>0.05) (Fig. 2e and f).

**Figure 2:**
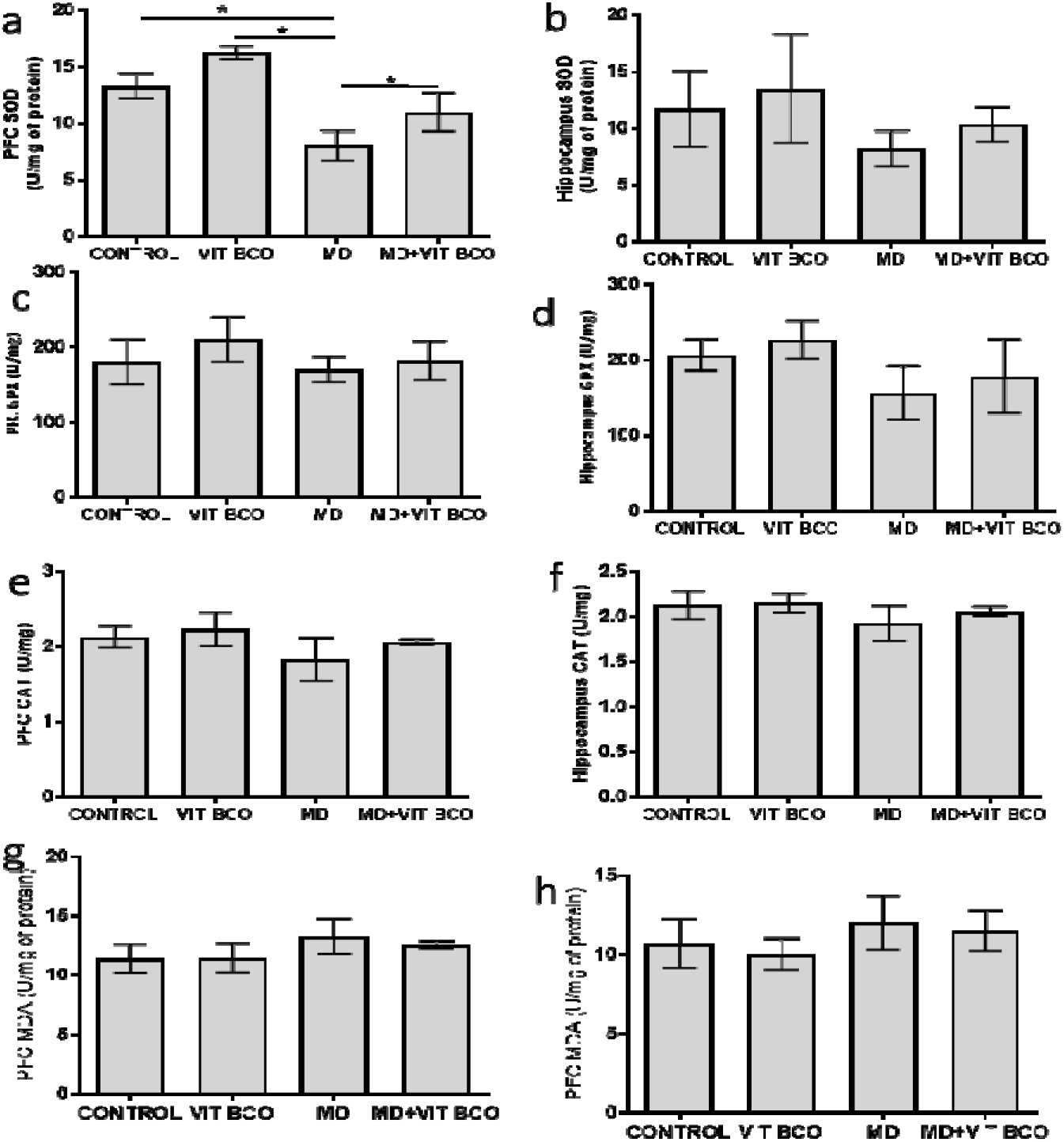
Neural antioxidant status and lipid peroxidation of rats. VIT BCO= Vitamin B complex, MD= maternal deprivation, MD + VIT BCO = maternal deprivation + Vitamin B complex. In Fig. a, SOD level in MD group was significantly reduced compared to experimental groups (p<0.05). Also in Fig. b, SOD level was reduced compared to control (p>0.05) and VIT BCO (p>0.05) group. In Fig. c, there was a decrease in GPx level compared to VIT BCO (p>0.05), but a slight decrease when compared to control (p>0.05) and MD+VIT BCO (p>0.05). Fig. d showed decrease in GPx level compared to control (p>0.05) and VIT BCO (p>0.05) group while slight changes existed between MD and MD+VIT but were not significant. Relative to the Control (Fig. **e**), CAT (catalase) level reduced in MD group compared to control and VIT BCO but not significant (p>0.05); changes in between Control and MD were very minimal and not significant (p>0.05). Also in Fig. f, there was a slight decrease in CAT level in MD compared to control (p>0.05) and VIT BCO (p>0.05) group while minimal changes between MD and MD+VIT BCO but not significant (p>0.05). Fig. g revealed that MDA levels was highest in the MD group compared to control (p>0.05) and VIT BCO (p>0.05) group, with a slight decrease in MD+VIT BCO (p>0.05) group. Fig. h revealed, MDA level was highest in the MD group compared to control VIT BCO but was not significant (p>0.05). The difference between MD and MD+VIT BCO (p>0.05) was minimal. * was the significant difference at p<0.05.

**Figure 3:**
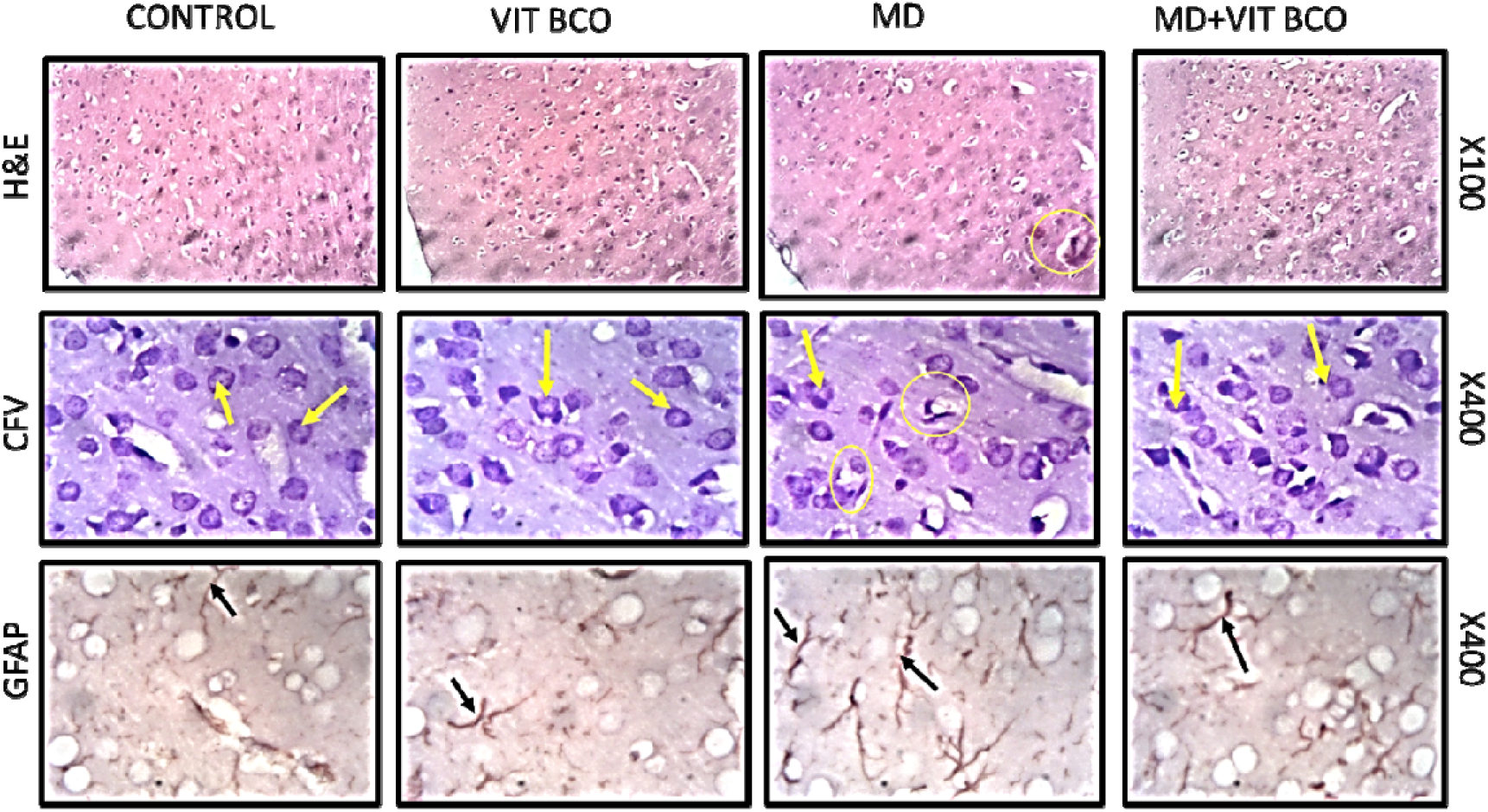
Representative photomicrographs of PFA-fixed prefrontal cortices of 35 days old rats stained with haematoxylin and eosin (H&E), cresyl fast violet (CFV) and anti-glial fibrillary acidic protein (GFAP) showing control, vitamin B complex-treated (VIT BCO), maternal deprivation (MD) and maternal deprivation + vitamin B complex-treated (MD+ VIT BCO). In H&E sections presented with characteristically stained large pyramidal dendrites with apical and basal dendrites jutting out of the soma (yellow arrows). The cells possessed intact neuropil with no sign of cellular fragmentation. Maternally deprived rats presented with reduced cellular density and fragmented neuropil as a sign of pyknosis (yellow circles). Maternally deprived rats treated with vitamin B complex presented with pyramidal cells slightly similar to the control. Nissl profile (CFV) showed that control and vitamin B complex had intensive chromatogenic pyramidal cells with prominent neuronal processes projecting out from the cell body (yellow arrow). The apical and basal dendrites projecting out of these cells also appeared to be chromatogenic as they were also positively stained for Nissl substances. However, maternally deprived rats presented with chromatolytic and pyknotic pyramidal cells indicated by poorly stained cell bodies (yellow circle). MD + vitamin B complex presented with mild chromatolytic changes (yellow arrow). Immunohistochemical staining with anti-GFAP showed that control and vitamin B complex had normal astrocyte distribution with no activation around the neuronal cells and generally low astrocytic densities (black arrows). Maternally deprived rats showed hypertrophied astrocytes and more number of astrocytes (red arrows) compared with the vitamin B complex group, and the deprived rats that received vitamin B complex.

**Figure 4:**
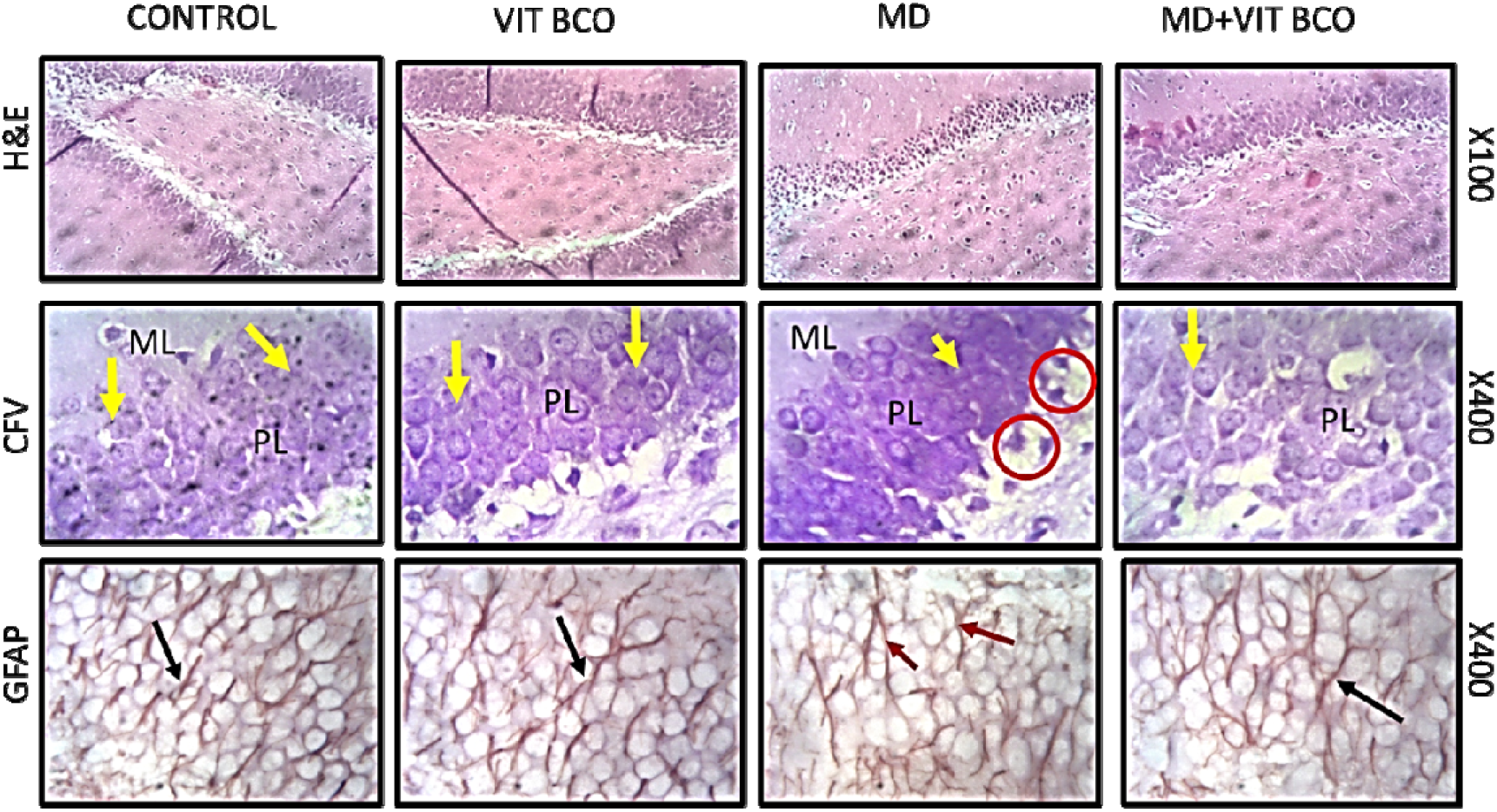
Representative photomicrographs of PFA-fixed hippocampi of 35 days old rats stained with haematoxylin and eosin (H&E), cresyl fast violet (CFV) and anti-glial fibrillary acidic protein (GFAP) showing control, vitamin B complex-treated (VIT BCO), maternal deprivation (MD) and maternal deprivation + vitamin B complex-treated (MD+VIT BCO). H&E sections revealed normal hippocampal morphology characterized by a distinct arrangement of pyramidal neurons in CA3, normal cell bodies with dendritic and axonal processes, as well as highly chromatogenic pyramidal cells and neuroglia (CFV). The granular layers and the molecular layer have a well-stained intensity and are well arranged. Maternal deprivation caused chromatolysis in pyramidal cells and neuroglia within the CA3 layers (red circle). Also, the layer of the granular and molecular cells appeared lightly stained. Treatment with vitamin B complex following maternal deprivation showed an improved chromatogenic property of the pyramidal cells and neuroglia within the CA3 when compared to maternally deprived rats. GFAP immunopositive cells (black arrows) in control and vitamin B complex treated group appeared sparse around neurons and between layers within the CA3, with regular processes, distribution and sizes within the neuropil. In both groups, astrocytic processes, cellular distribution and size were normal. However, there was increased astrocytic density with reactive astroglia in the granule cell layer of the maternally deprived rat. Maternal deprivation + vitamin B complex group showed improved astrocytic morphology and distribution, which had close similarities to those in the control group.

Cortico-hippocampal levels of malondialdehyde (MDA), a by-product of lipid peroxidation was elucidated. We found that the MD group presented with a slightly raised level of MDA in both brain region (p>0.05) when compared to the control and VIT BCO group (Fig. 2g and h). Cumulatively, MD had a differential affectation on the antioxidant defence system which culminated in an increased rate of lipid peroxidation. MD rats treated with VIT BCO manifested a cortical and hippocampal level of GPx, catalase and SOD that were similar to that of the control group. VIT BCO play a neuroprotective role by sustaining the level of MDA in the PFC and hippocampus to a level comparable to that of the control.

### 3.4 Histomorphology, Nissl profile and astrocyte expression in the prefrontal cortex and hippocampus

Histological presentation of the prefrontal cortex showed the external pyramidal layer of control and vitamin B complex-treated rats presented with typical cellular density and proper cortical delineation of cells. The pyramidal cells had large cell bodies with adjoining neuronal processes and supporting cells in the neuropil; soma and dendritic processes were properly stained and distributed, and there were intact Nissl bodies within the cell bodies. Maternally deprived rats presented with reduced cellular density, poorly stained pyramidal cells with fragmented neuropils as signs of pyknosis; mild chromatolysis and poorly stained cells of external pyramidal layers were also present. Rats that received vitamin B complex treatment after maternal deprivation had a histological presentation similar to those in control.

The *Cornu Ammonis* 3 (CA3) region of the hippocampus of the control and vitamin B complex-treated rats had large pyramidal cells with normal cell bodies, dendritic and axonal processes that were well expressed across hippocampal neuropil; the granular and molecular layers were also well stained with intact Nissl bodies in the cells. Maternally deprived rats revealed the presence of degenerating pyramidal cells, chromatolytic pyramidal cells and poorly stained Nissl bodies within the CA3 layer. The hippocampus of rats that received vitamin B complex intervention after deprivation showed characteristics that were similar to those of the control and vitamin B complex-treated rats.

The expression of astrocytes in the prefrontal cortex and hippocampus appeared normal as evidenced by the presence of sparse immunopositive cells in the control and vitamin B complex-treated brain. The rats that were maternally deprived presented with a slight increase in astrocytic density, which was again reduced in those that received vitamin B complex.

## 4 Discussion

The present study characterized the behavioural alterations, neurochemical perturbation, and histomorphological modifications in the PFC and hippocampus of maternally deprived experimental animals, detailing the possible neuroprotective roles of vitamin B complex supplementation. Furthermore, the changes in the body and brain weights were documented.

Maternally deprived rats have been reported to show a significant decrease in the body weight in adolescence and adulthood (Llorente et al., 2007). This finding is consistent with results in the present study, where we noted a significant reduction in the body weight of animals that were maternally deprived. The weight loss has been attributed to lack of milk ingestion during the critical period of development period resulting in an alteration in the developmental programming of hypothalamic energy regulatory circuits (Llorente et al., 2007).

Our interest in the hippocampus and prefrontal cortex is because earlier studies have reported an oxidative perturbation in these brain regions of MD rats (Diehl et al., 2012; Holland et al., 2014). Furthermore, the hippocampus is important in memory formation and consolidation (Izquierdo and Medina, 1997), and it is highly susceptible to oxidative damage (Plotsky et al., 2005). The prefrontal cortex, on the other hand, partakes in cognitive functions such as working memory and memory consolidation (Kesner and Churchwell, 2011). Early perinatal stress can cause various short- and long-term disturbances in behavioural performances (Koehl et al., 2001). Findings from the present study showed that maternal deprivation caused a significant reduction in the memory indices of the experimental animals while increasing anxiety-like behaviours in the said animals. These inferences were drawn from the increase in escape latency period in the Morris water maze, reduction in the percentage correct alternation in the Y-maze and unwillingness of the maternally deprived rats to explore the open field test. Earlier work by, Oitzl *et al*. (2000) reported that a single 24-h maternal deprivation led to impairment in spatial learning ability in the Morris water-maze test (Oitzl et al., 2000). However, a tendency for improvement in water-maze learning in maternally deprived adult rats has also been reported (Enthoven et al., 2008). Breast milk, unlike formula feeds, comprises long-chain polyunsaturated fatty acids, like docosahexaenoic and arachidonic acids which are vital in neurodevelopment (Amore et al., 2003). Missing these nutrients in addition to early life stressful events (such as maternal separation) have been reported to affect normal brain development as well as implicated in the psychopathology of behavioural disorders such as anxiety, depression, schizophrenia and memory decline (Amore et al., 2003; Takase et al., 2012; Zugno et al., 2013). Intervention with vitamin B complex in this study resulted in enhanced behavioural performances in the memory behavioural paradigms.

Maternal deprivation is one of the most compelling natural stressors during the early stages of development (Neves et al., 2015; Uysal et al., 2005). Events from this type of stress can result in permanent deficits during adulthood (Neves et al., 2015). Activities of antioxidant enzymes and lipid peroxidation in the frontal cortex and hippocampus were quantified in the present study. Superoxide dismutase catalyzes the conversion of highly reactive superoxide to less reactive hydrogen peroxide. Catalase is the main enzyme responsible for the removal of hydrogen peroxide from normal tissues. By breaking down hydrogen peroxide, this enzyme does not cause the generation of any other reactive oxygen species. Findings from this study showed that maternal deprivation caused a reduction in superoxide dismutase activities in both brain regions. This finding is not in conformity with what was reported by Markovic *et al*. (2017) that maternal deprivation brought about an increase in the activity of superoxide dismutase in the prefrontal cortex and hippocampus (Markovic *et al*., 2017). This report suggests the role that maternal separation during the critical period of neurodevelopment plays in perturbing the antioxidant defence system later in life. This kind of perturbation will result in the inability of the brain to adequately clear intrinsically produced reactive species, thereby initiating a cascade of chemical events that will exacerbate lipid peroxidation. Interestingly, malondialdehyde (an end product of lipid peroxidation) was not significantly altered in the maternal deprivation group. Findings from this study showed that vitamin B complex conferred a degree of protection on the neural activities in the PFC and hippocampus in the maternally deprived group. We noted that animals treated with vitamin B complex presented with normal levels of antioxidant enzymes with no changes in their hippocampal and cortical lipid peroxidation profile. Therefore, enhancing the intrinsic antioxidant defence system is an interesting way to restore the neural activities in maternally deprived animals to normal. A previous study has reported that boosting the antioxidant status of the brain in deprived animals through physical activities helped to prevent behavioural decline and oxidative stress that comes with maternal deprivation (Neves et al., 2015). It was demonstrated that physical exercise prevents oxidative damage (lipid peroxidation) in the hippocampus and prefrontal cortex (Neves et al., 2015).

The effect of MD during the critical period of development is connected to the neural changes that happen during this period. For instance, the majority of granular neurons of the hippocampus develop and extend their axons between postnatal day 1 and 21 (Amaral and Dent, 1981). This fact explains the histomorphological alterations observed in the present study. Even though the general histology did not show much difference across the experimental groups, maternally deprived rats presented with increase chromatolysis and astrocyte activation in the PFC and hippocampus. Increased astrocyte expression is an indicator of MD-enhanced neuroinflammation (Gracia-Rubio et al., 2016; Réus et al., 2017). Vitamin B complex treatment prevented excessive chromatolysis and neuroinflammation in this study. Maternal deprivation caused astrogliosis, which was observed as increased astrocytic densities, and cytoplasmic vacuoles within the hippocampal layer and prefrontal cortex. This result is in agreement with the study that showed that maternal deprivation modulates adult GFAP levels in hippocampus, cerebellum and cortex (Llorente et al., 2009; López-Gallardo et al., 2008; Omotoso et al., 2020). These findings elucidate the neuroprotective properties of vitamin B complex in preventing astrogliosis, as well as oxidative stress in the hippocampus and prefrontal cortex of maternally deprived rats.

## 5 Conclusion

Cumulatively, we have shown that Vitamin B complex significantly counteracted maternal deprivation-induced neurotoxicity by eliciting a significant protective effect on neurobehaviour, boosting antioxidant defence system and sustaining the integrity of neuronal cells in the hippocampus and prefrontal cortex.

## Notes

### Competing Interest Statement

The authors have declared no competing interest.

